# A novel micronemal protein, Scot1, is essential for apicoplast biogenesis and liver stage development in *Plasmodium berghei*

**DOI:** 10.1101/2024.04.23.590848

**Authors:** Ankit Ghosh, Akancha Mishra, Raksha Devi, Sunil Kumar Narwal, Nirdosh, Pratik Narain Srivastava, Satish Mishra

**Affiliations:** Division of Molecular Microbiology and Immunology, CSIR-Central Drug Research Institute, Lucknow 226031, India; Academy of Scientific and Innovative Research (AcSIR), Ghaziabad 201002, India

**Keywords:** GAP, *Plasmodium*, microneme, liver stage, malaria, preerythrocytic, protection, sporozoite, vaccine

## Abstract

*Plasmodium* sporozoites invade hepatocytes, transform into liver stages, and replicate into thousands of merozoites that infect erythrocytes and cause malaria. Proteins secreted from micronemes play an essential role in hepatocyte invasion, and unneeded micronemes are subsequently discarded for replication. The liver-stage parasites are potent immunogens that prevent malarial infection. Late liver stage-arresting genetically attenuated parasites (GAPs) exhibit greater protective efficacy than early GAP. However, the number of late liver-stage GAPs for generating GAPs with multiple gene deletions is limited. Here, we identified Scot1 (Sporozoite Conserved Orthologous Transcript 1), which was previously shown to be upregulated in sporozoites, and by endogenous tagging with mCherry, we demonstrated that it is expressed in the sporozoite and liver stages in micronemes. Using targeted gene deletion in *Plasmodium berghei*, we showed that Scot1 is essential for late liver-stage development. *Scot1* KO sporozoites grew normally into liver stages but failed to initiate blood-stage infection in mice due to impaired apicoplast biogenesis and merozoite formation. Bioinformatic studies suggested that Scot1 is a metal/small molecule carrier protein. Remarkably, supplementation with metals in the culture of infected *Scot1* KO cells did not rescue their phenotype. Immunization with *Scot1* KO sporozoites in C57BL/6 mice confers protection against a malaria challenge via infection. These proof-of-concept studies will enable the generation of *P. falciparum Scot1* mutants that could be exploited to generate GAP malaria vaccines.

**Importance:** Malaria parasites experience significant bottlenecks as transmitted to the mammalian host during a mosquito bite. Sporozoites invade liver cells, reproducing into thousands of merozoites, which are released after liver cell ruptures. The specific arrest of sporozoites during liver stage development acts as a powerful immunogen and provides sterile protection against sporozoite infection. GAP leading to an arrest in late liver stage development offers superior protection. Here, we report that a micronemal protein, Scot1, is essential for parasite maturation in the liver. Deletion of Scot1 resulted in impaired apicoplast biogenesis and merozoite formation. Vaccination with *Scot1* KO sporozoites protects against malaria challenge. We have identified a late arresting GAP that will aid in developing new as well as safeguarding existing whole parasite vaccines.

## Introduction

Malaria has a tremendous negative impact on human health and the economy, a trend that unfortunately continues even today. Malaria-causing *Plasmodium* parasites are responsible for more than 249 million cases and 0.61 million deaths in 2022 (1). *Plasmodium* sporozoites are transmitted via the bite of an infected *Anopheles* mosquito, which invades the liver and transforms into exoerythrocytic forms (EEFs). Within hepatocytes, the parasite undergoes intracellular replication and forms thousands of merozoites, which initiate pathogenic erythrocytic stages (2). The parasite must eliminate unnecessary micronemes throughout this development process to create room for newly synthesized stage-specific organelles and proteins (3). It was recently demonstrated that the *Plasmodium* autophagy pathway is essential for eliminating unneeded micronemes during the development of the parasite liver stages (4). However, the contributions of micronemal resident proteins to eliminating micronemes remain unknown. To date, micronemal proteins are secreted during invasion and play a crucial role in the gliding motility and infectivity of *Plasmodium* sporozoites (5).

The micronemal protein TRAP (Thrombospondin-related anonymous protein) links the sporozoite actin/myosin motor to the extracellular substrate and was shown to be essential for gliding motility (5, 6). Other micronemal proteins, such as SPECT (sporozoite microneme protein essential for cell traversal), MAEBL (merozoite apical erythrocyte-binding ligand), P36, P52, GAMA (GPI-anchored micronemal antigen), CelTOS (cell traversal for ookinetes and sporozoites), TRP1 (Thrombospondin-related protein 1), S6/TREP and AMA1 (Apical Membrane Antigen 1), have been characterized in *Plasmodium* (5). CelTOS, MAEBL, SPECT, and AMA1 are involved in cell traversal (5, 7). In addition to erythrocyte invasion, AMA1 was shown to be secreted from sporozoites and drive their invasion (5, 8). S6/TREP was found to be important for parasite motility and efficient malaria transmission (9). P36 and P52 form a complex and play an essential role in the productive invasion of sporozoites (5). Disruption of P52 and P36 leads to the attenuation of parasites in the liver and confers sterile immunity against infection (10). GAMA and TRP1 were found to be important for sporozoite egress from oocysts (11, 12).

The emergence and spread of artemisinin-resistant *P. falciparum* threatens the control and elimination of malaria. New drugs and a highly effective malaria vaccine are urgently needed. The World Health Organization (WHO) has recommended using the circumsporozoite protein (CSP)-based RTS,S vaccine despite its modest efficacy (13). R21/Matrix-M is the second malaria vaccine recommended by the WHO (14). The major drawback of this recombinant vaccine is the lack of an efficient and durable immune response, which may not be suitable for long-term usage. Occasionally, subunit vaccines do not elicit as strong or long-lasting immune response as whole-organism vaccines. The radiation-attenuated sporozoite (RAS) vaccine has existed for several decades and has been proven to have long-term protective effects (15). Moreover, the induction of sterile immunity by the RAS was very encouraging. Nonetheless, one of the main drawbacks of this approach is that either underirradiation leads to breakthrough infection, or overirradiation of sporozoites leads to failure in initiating optimal preerythrocytic immunity (16). Immunization with live sporozoites attenuated by genetic modification has attracted much attention because they have been shown to produce protective immune responses equal to or even greater than those produced by irradiated sporozoites in rodent models (17). The protection of these attenuated sporozoite vaccines involves antibodies elicited against sporozoite antigens that neutralize their ability to invade hepatocytes (18). Moreover, this protection is mediated through CD8+ T-cell responses that target infected hepatocytes (19).

The *P. falciparum* RAS vaccine expresses thousands of proteins and elicits superior protection compared to RTS,S; however, it does not confer complete protection in endemic areas (20) and requires improvement. Recent advances in *Plasmodium* genetics have enabled the generation of many GAPs, which have overcome the limitations of using RAS as a whole-organism vaccine. However, except for a few GAPs, most of these GAPs are blocked at the early to mid-liver stage, which limits the antigen breadth and biomass for superior immune protection (21). It was shown that immunization with *P. falciparum* sporozoites under a drug cover allows liver-stage parasites to mature, generating durable protection at lower doses than the *P. falciparum* RAS sporozoite vaccine (22). Although arrested parasites are a source of antigen for effective immune system priming, their antigenic repertoire induces cross-stage immunity only when parasites are blocked at the late liver stage (23). Since late liver stages exhibit a subset of antigens common to blood stages, identifying sporozoite genes that can yield a late arrest mutant will have a broader impact on developing an efficacious GAP vaccine. Here, we disrupted the function of the *P. berghei* micronemal protein Scot1 and investigated its role during late liver stage development. Immunization with late arresting GAP confers protection against infectious sporozoites.

## Results

### Scot1 is a highly conserved *Plasmodium*-specific protein

Phylogenetic analysis revealed that Scot1 is a *Plasmodium*-specific protein that is absent in other organisms (Figure S1). Scot1 was found to be highly conserved among *Plasmodium* species (Figure S2A). The identity matrix showed 100% sequence similarity between *P. berghei*, *P. yoelii* and *P.vinckei* and 92.1% similarity between *P. berghei* and *P. falciparum* and 90.9% similarity with *P. knowlesi* (Figure S2B). The Scot1 protein lacks a signal sequence and transmembrane domain.

### Scot1 is expressed during the sporozoite and liver stages and is localized to the microneme

To study the expression and localization of Scot1, the gene was endogenously tagged with 3XHA-mCherry using double crossover homologous recombination (Figure S3A). Correct integration of the *Scot1* gene was confirmed by diagnostic PCR (Figure S3B). The development of transgenic parasites was analysed in mosquito and mammalian hosts. We found that the C-terminal tag did not affect parasite development throughout the life cycle (Figure S3C and S3D). The Scot1 promoter expressed the Scot1-mCherry protein in oocyst and salivary gland sporozoites and liver stages at 12, 24, and 62 hpi (Figure 1A, B and S3E). mCherry expression was not detected in the blood, gamete, ookinete, or liver stages at 36 or 48 hpi. The expression of the Scot1-3XHA-mCherry fusion protein was also confirmed by immunoblotting using an anti-mCherry antibody (Figure 1C). We then analysed the localization of Scot1-3XHA-mCherry in sporozoites and liver stages. Sporozoites were stained with anti-mCherry and anti-TRAP antibodies. We observed a granular localization pattern of Scot1-3XHA-mCherry, which colocalized with the TRAP signal in sporozoites, indicating that Scot1 is a micronemal protein (Figure 1D). These results indicate that Scot1 is expressed in sporozoites and liver stages and is localized to the microneme.

**Figure 1.**
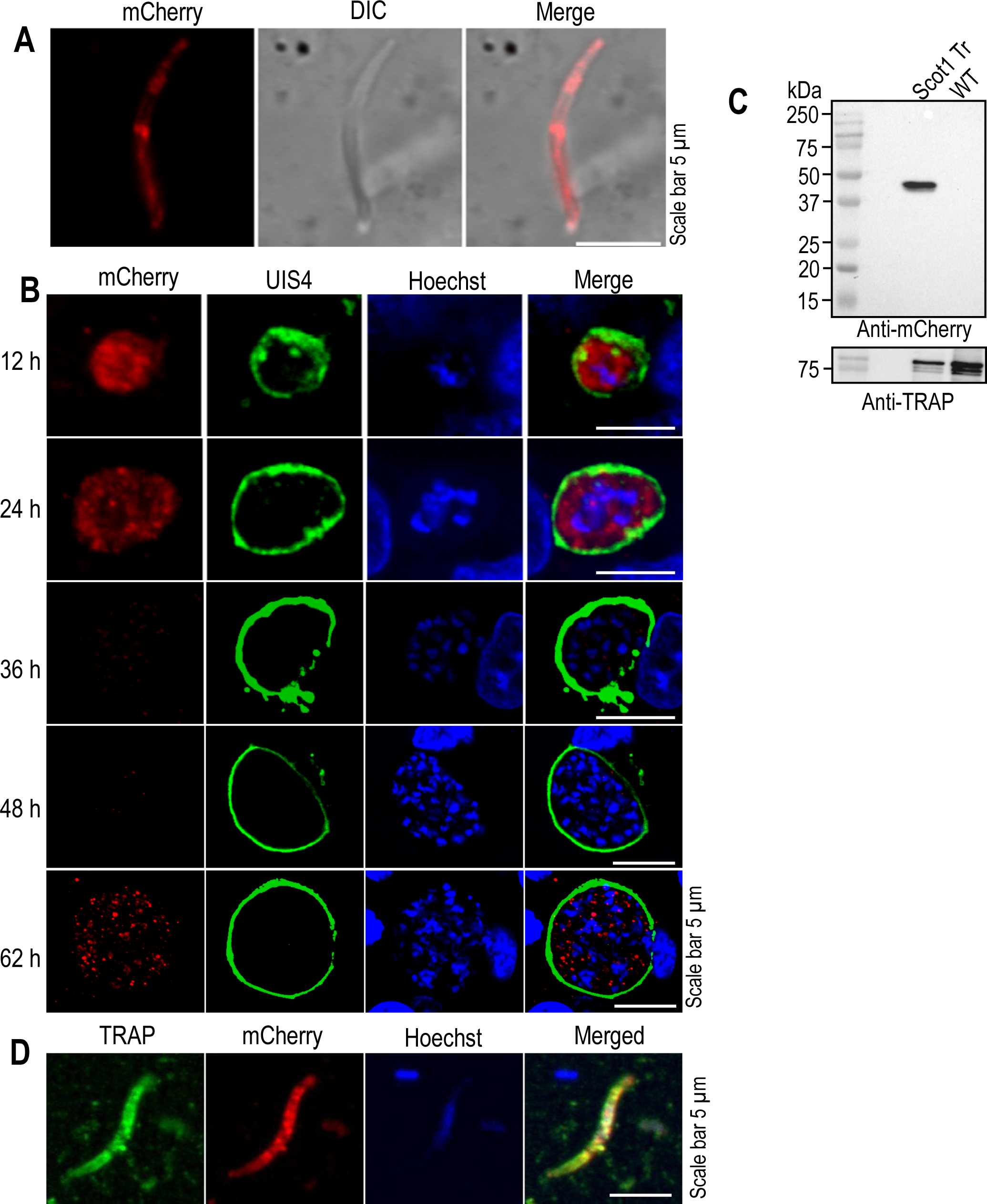
Expression and localization of Scot1 in *P. berghei*. **(A)** Scot1-3XHA-mCherry salivary gland sporozoites showing mCherry expression pattern. **(B)** HepG2 cells infected with Scot1-3XHA-mCherry sporozoites were harvested at different time points. Cultures harvested at 12, 24, 36, 48, and 62 hpi were stained with mCherry and UIS4 antibodies, and nuclei were stained with Hoechst. mCherry expression was observed at 12, 24, and 62 hpi but at 36 and 48 hpi. **(C)** Confirmation of the expression of the Scot1 fusion protein in the Scott1-3XHA-mCherry transgenic parasites using an anti-mCherry antibody. A 39 kDa band was detected in the lysate of the Scot1 transgenic sporozoites but not in the lysate of the WT parasites. The blot was stripped and reprobed with an anti-TRAP antibody as a loading control. **(D)** IFA of Scot1-3XHA-mCherry transgenic sporozoites using anti-TRAP and anti-mCherry antibodies. TRAP and mCherry were colocalized (PCC=0.9 of 10 sporozoites).

### Scot1 is dispensable in the *P. berghei* blood and mosquito stages

To investigate the role of Scot1 in the parasite life cycle, we disrupted the gene in *P. berghei* using double crossover homologous recombination (Figure S4A). Resistant GFP (green fluorescent protein)-expressing parasites were confirmed by fluorescence microscopy (Figure S4B), and correct genomic integration was confirmed by diagnostic PCR (Figure S4C). A Scot1-complemented parasite line was generated by transfecting the *Scot1* expression cassette into *Scot1* KO schizonts (Figure S4D). Restoration of the *Scot1* locus was confirmed by diagnostic PCR (Figure S4E). To determine whether the deletion of *Scot*1 affected the dynamics of blood growth, two groups of mice were intravenously inoculated with WT GFP and *Scot1* KO parasites. Parasite growth was monitored by making Giemsa-stained blood smears. No difference was observed between WT GFP and *Scot1* KO parasites (Figure S4F). For the phenotypic characterization of the *Scot1* KO parasites, the mosquito cycle was initiated by infecting *A. stephensi* mosquitos with *Scot1* KO or WT GFP parasites. On day 14 postinfection, the mosquito midgut was dissected to check for the presence of oocysts, which was comparable in both WT GFP and *Scot1* KO parasites (Figure S5A and B). The sporogony in oocyst and sporozoite numbers were also normal (Figure S5C and D). Furthermore, on days 18-22 after a blood meal, the salivary glands were observed under a fluorescence microscope, and the sporozoite numbers were counted, which revealed a normal sporozoite load and number (Figure S5E and F). These results demonstrate that the deletion of *Scot*1 does not affect parasite development in the blood or mosquito stages.

### *Scot1* KO sporozoites infect the liver but fail to initiate blood-stage infection in mice

To assess the in vivo infectivity of the *Scot1* KO parasites, salivary gland sporozoites were injected intravenously into C57BL/6 mice or infected by mosquito bites, and the appearance of the parasites in the blood was monitored by making Giemsa-stained blood smears. The WT GFP-inoculated mice were positive on day 3 post infection, whereas *Scot1* KO sporozoites failed to initiate blood-stage infection (Table 1). Scot1 gene complementation restored the KO phenotype (Table 1), indicating the specificity of the gene function. Micronemal proteins were previously shown to be required for the invasion of hepatocytes (24, 25). Therefore, we checked the invasion ability of the *Scot*1 KO sporozoites and found that they were normal (Figure S6). To analyse the progression of the parasites in vivo, the livers of infected mice were harvested at 40 and 55 hpi, and the parasite burden was quantified by amplifying 18S rRNA using real-time PCR. We found no difference in the 18S rRNA copy number at 40 hpi, but it was significantly lower at 55 hpi in the *Scot1* KO parasites than in the WT GFP parasites (Figure 2). These results provide evidence that *Scot1* KO parasites grow normally until the mid- to late-liver stage but do not mature and fail to initiate blood-stage infection in mice.

**Figure 2.**
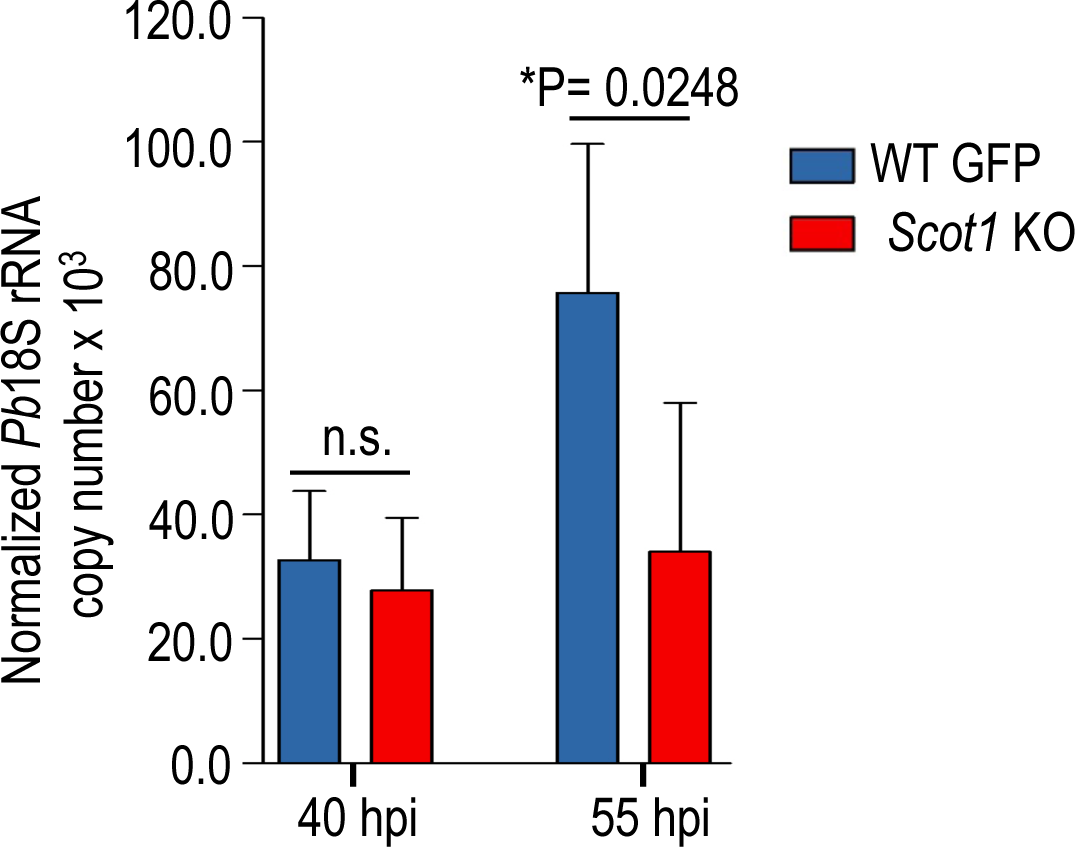
*Scot1* KO parasites exhibit major defects during late liver stage development. (**A**) To quantify the parasite burden, infected mouse livers were harvested at 40 and 55 hpi, RNA was isolated, and transcripts were quantified using real-time PCR. The *P. berghei* 18S rRNA copy number was comparable in the WT GFP and *Scot*1 KO parasites at 40 hpi (P=0.5233, Student’s t test) but decreased in the *Scot*1 KO parasites compared to the WT GFP parasites at 55 hpi (*P=0.0248, Student’s t test). The data are presented as the means ± SDs; n = 5 mice per group.

**Table 1.**
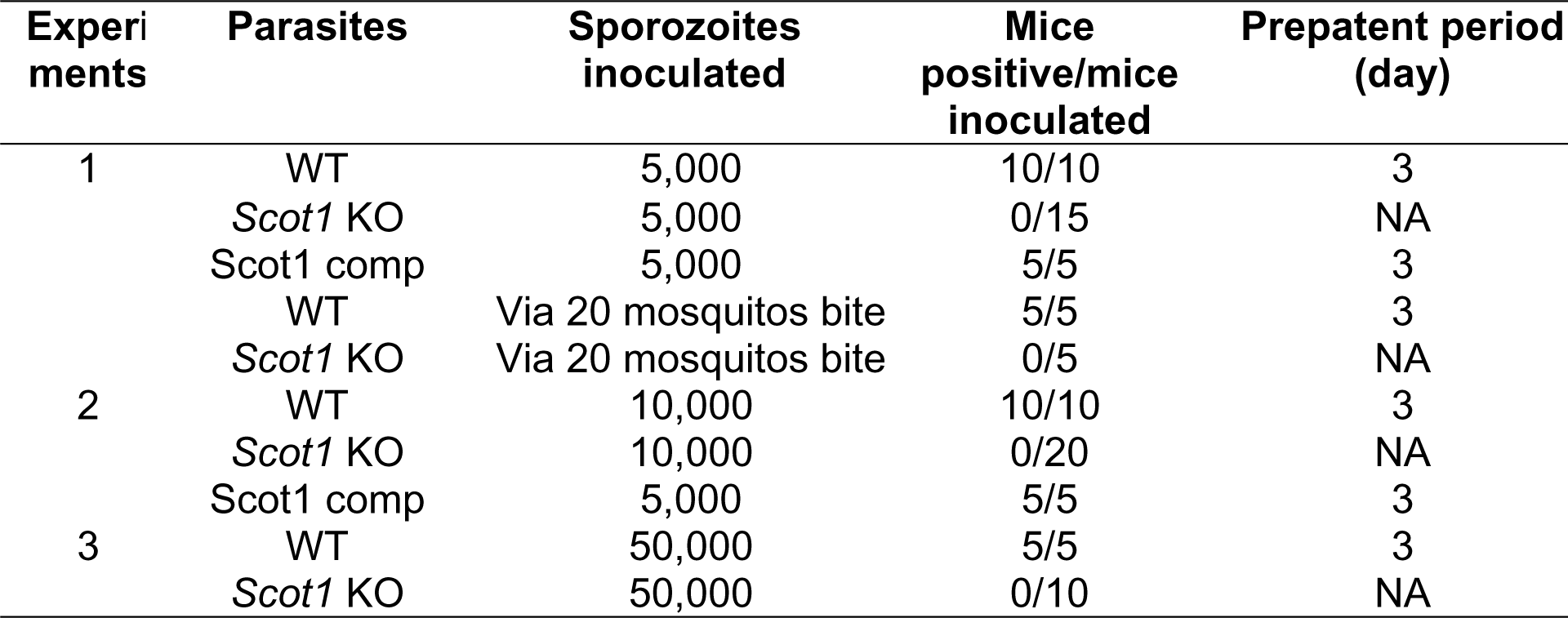
Infectivity of salivary gland sporozoites in C57BL/6 mice. Blood smears were examined daily from day 3 onwards, and the mice were considered negative if parasites were not detected by day 20. NA, not applicable.

### *Scot1* KO EEFs grow normally in size

The normal invasion and failure of *Scot1* KO sporozoites to initiate blood-stage infection in mice suggest that either they failed to develop into EEFs or egress from hepatocytes. To further investigate the liver stage development of the *Scot1* KO parasites, HepG2 cells were infected with sporozoites and fixed at different time points. The liver stages of the WT GFP and *Scot1* KO parasites showed similar growth, EEF numbers, and sizes (Figure 3A-C). *Scot1* KO EEF growth, number, and size analysis revealed no apparent aberrant phenotype; however, there was evidence of reduced nuclear division at 62 hpi. Next, we observed the culture for the formation of detached cells at 62 hpi. We found detached cells in the WT GFP culture but not in the *Scot1* KO-infected culture (Figure 3D). Next, we checked the infectivity of detached cells from WT GFP and culture supernatant from *Scot1* KO parasites in Swiss mice. We found infection in mice injected with WT GFP-detached cells; no infection was observed in the KO-injected group (Table 2).

**Figure 3.**
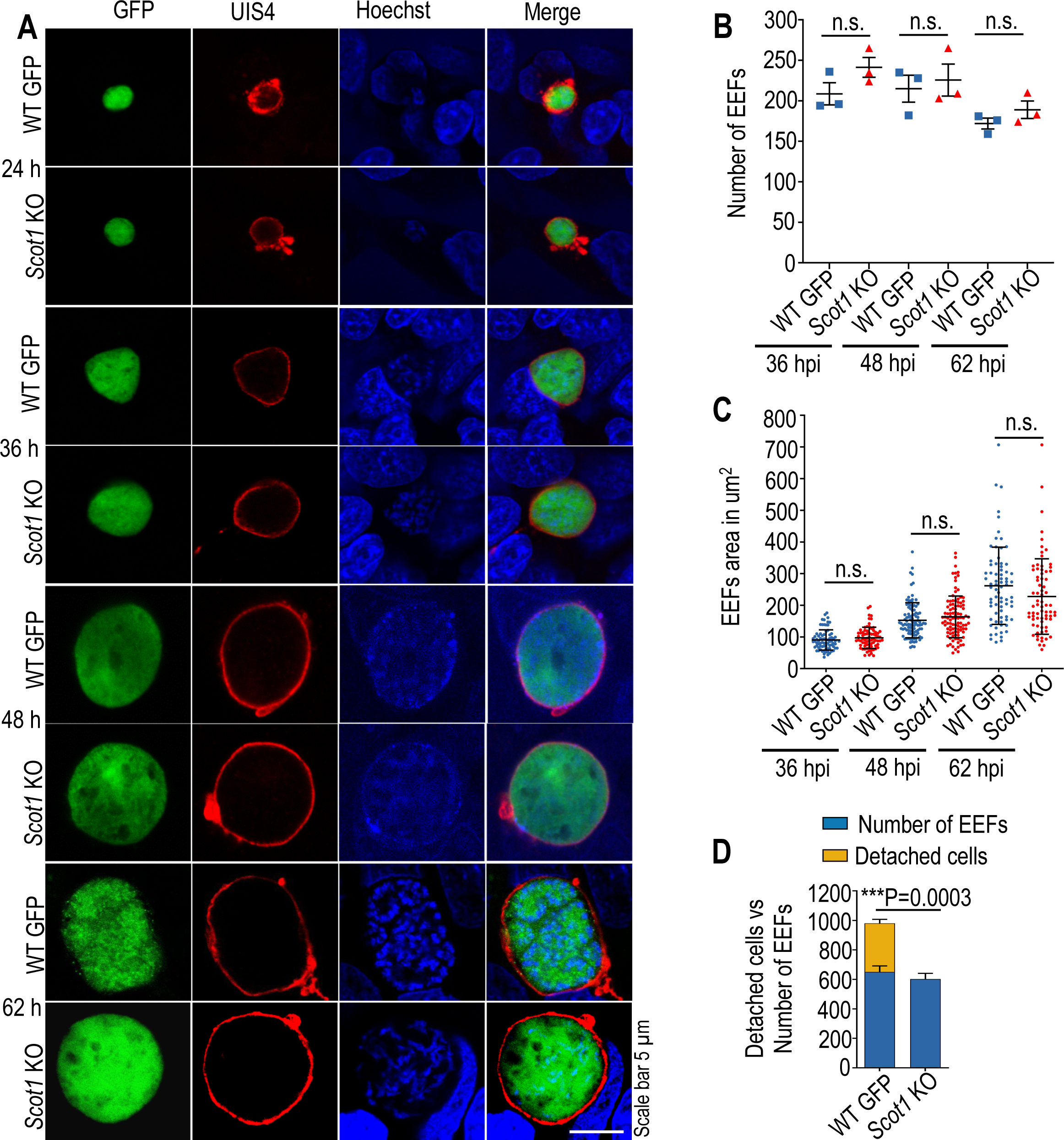
*Scot1* KO EEFs exhibit normal growth but fail to mature into hepatic merozoites. **(A)** HepG2 cells infected with the *Scot*1 KO or WT GFP sporozoites were harvested at different time points. Cultures harvested at 24, 36, 48, and 62 hpi were stained with anti-UIS4 antibody, and host and parasite nuclei were stained with Hoechst. The EEFs grew normally until 48 hpi and showed impaired nuclear division at 62 hpi. **(B)** The number of EEFs in the *Scot1* KO parasites at 36 (P=0.1426), 48 (P=0.7006) and 62 hpi (P=0.2518) was not significantly different from that in the WT GFP. The data were obtained from three independent experiments performed in duplicate and are presented as the mean ± SEM. **(C)** The EEF area at 36 (P=0.2064), 48 (P=0.2084) and 62 hpi (P=0.0908) was comparable between the WT GFP and *Scot1* KO parasites. The data were obtained from three independent experiments performed in duplicate and are presented as the mean ± SEM. **(D)** The number of EEFs from the *Scot1* KO and WT GFP parasites. Despite a comparable number of EEFs, *Scot1* KO parasites failed to release detached cells into the culture supernatant (***P<0.0003). The data were obtained from a single experiment performed in duplicate and are presented as the mean ± SD. Student’s t test was used to determine the statistical significance.

**Table 2.** Infectivity of the merosomes in Swiss mice. Blood smears were examined daily from day 1 onwards, and the mice were considered negative if parasites were not detected by day 20. NA, not applicable.

| Experiment | Parasite | Number of detached cells injected | Mice positive/mice inoculated | Prepatent period (day) |
| --- | --- | --- | --- | --- |
| 1 | WT | 10 | 5/5 | 2.4 |
|  | *Scot1* KO | Supernatant | 0/5 | NA |
| 2 | WT | 100 | 5/5 | 1.2 |
|  | *Scot1* KO | Supernatant | 0/5 | NA |

### *Scot1* KO parasites exhibit impaired apicoplast biogenesis and fail to mature into hepatic merozoites

Next, we analysed late liver-stage parasites by immunostaining with ACP (Acyl Carrier Protein) and MSP1 (Merozoite Surface Protein 1) antibodies. Loss of *Scot1* severely compromised apicoplast biogenesis, resulting in a consequent loss of MSP1 staining and no merozoite formation (Figure 4A and B). We found impaired nuclear division and a significant decrease in the number of nuclei in the *Scot1* KO parasites compared to those in the WT GFP parasites (Figure 4C). These results indicated the role of Scot1 during late liver-stage development.

**Figure 4.**
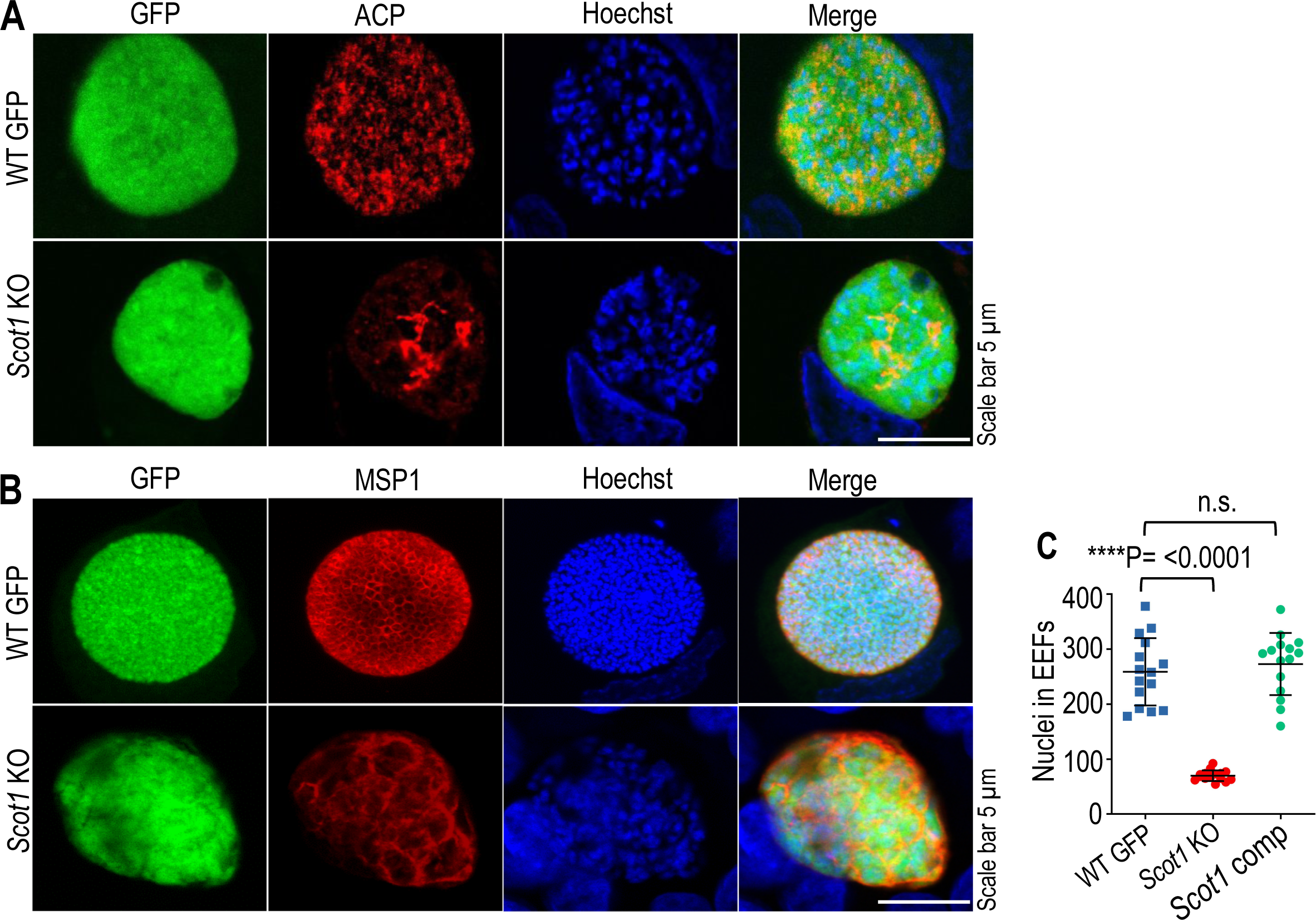
*Scot1* KO EEFs exhibit impaired late liver stage development. **(A)** Culture fixed at 62 hpi was immunostained with the apicoplast marker anti-ACP antibody. WT GFP and *Scot1* KO parasites showing apicoplast branching patterns. (**A**) Infected cultures harvested at 62 hpi were immunostained with an anti-MSP1 antibody to visualize the development of hepatic merozoites, and DNA was stained with Hoechst. No merozoites were observed in the *Scot1* KO parasites. We found normal segregation of nuclei and the formation of merozoites in the WT GFP parasites but not in the *Scot1* KO parasites. **(B)** Nuclei were counted in WT GFP, *Scot1* KO and Scot1 comp EEFs at 62 hpi, and a significant decrease in nuclear count was observed in *Scot1* KO parasites (****P<0.0001, Student’s t test). No significant difference was observed between WT GFP and Scot1 comp parasites (P= 0.5166, Student’s t test).

### Scot1 is not required for the elimination of micronemes

After sporozoite invasion, unneeded micronemes are expelled into the PV during liver stage development (3, 26). To determine whether the microneme-localized protein Scot1 is involved in eliminating micronemes, we monitored its distribution pattern during liver stage development using an anti-TRAP antibody. We found that by 24 hpi, the micronemes were directed toward the PV membrane of the parasite, and by 40-55 hpi, all the micronemes were expelled into the PV membrane, indicating normal elimination of the micronemes from the *Scot1* KO and WT GFP parasites (Figure 5). These data indicate that Scot1 is dispensable for the elimination of micronemes.

**Figure 5.**
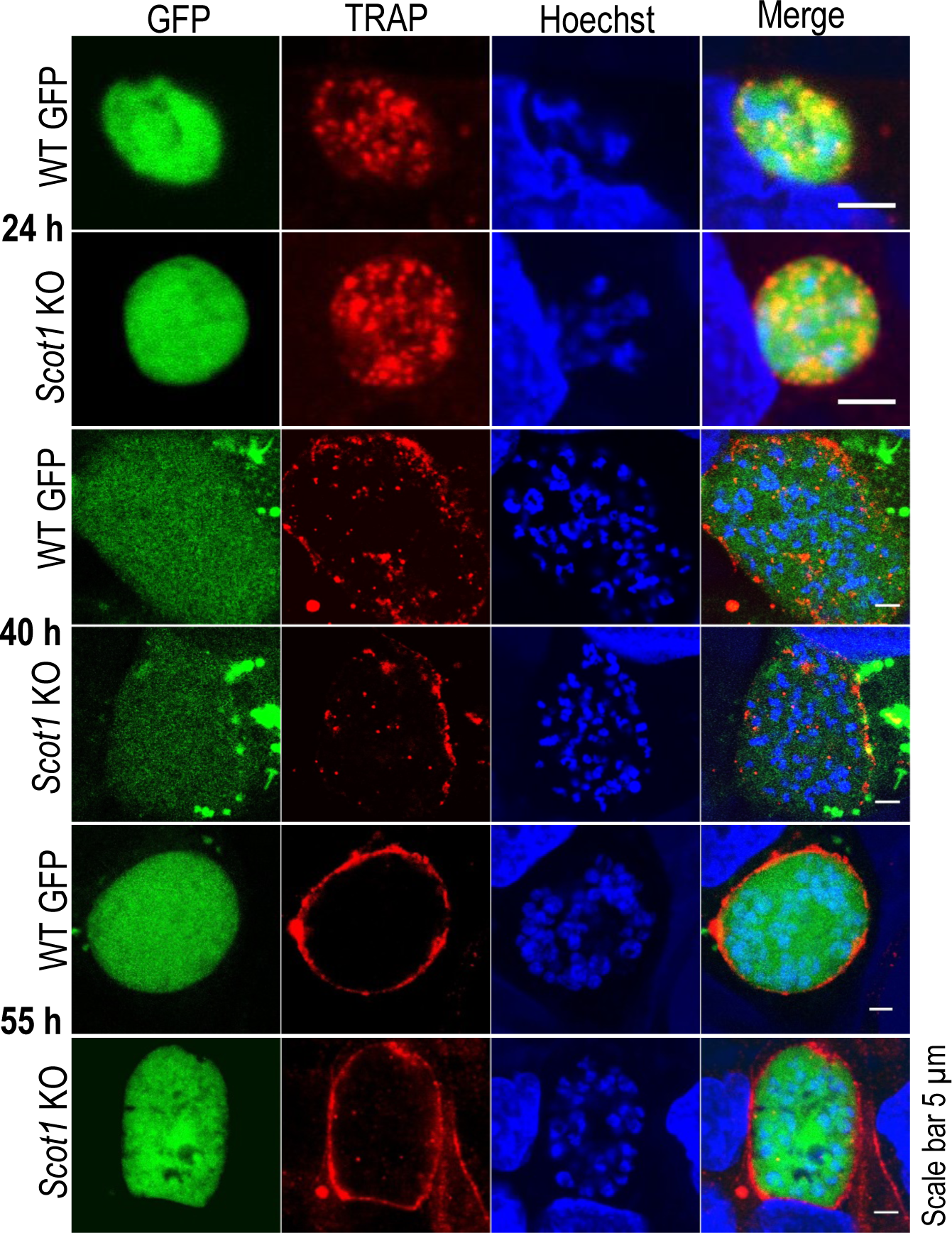
The *Scot1* KO parasites expel micronemes normally. HepG2 cultures infected with the *Scot1* KO or WT GFP parasites were fixed at 24, 40, and 55 hpi and immunostained with anti-TRAP antibodies. During the liver stage of *Plasmodium* development, micronemes were expelled into the PV in both the *Scot1* KO and WT GFP parasites. Nuclei were stained with Hoechst.

### Bioinformatic studies suggest that Scot1 is a metal/small molecule carrier protein

Structural modelling of PfScot1 revealed a beta-pleated sheet structure twisted in itself, forming an open barrel shape (Figure 6A). Structural superposition with the SCOP and PDB databases using the FATCAT and DALI web services and subsequent superposition of the results in UCSF Chimera showed that the model aligns with the haemoglobin linker chain L1 (chain M of PDB ID 2GTL (red)) with a 0.9 Angstrom RMSD value (Figure 6B) and chain A of the crystal structure of the vaccine antigen Transferrin Binding Protein B (TbpB) (PDB ID: 4O4X (red)), a metal transport protein with a 0.8 Angstrom RMSD value (Figure 6C). The top hits from the structure-based phylogenetic analysis further emphasize the possible function of PfScot1 as a carrier of metal ions or small molecules (Figure 6D). Proteins with the closest structural similarity to that of Scot1 are listed in Table S1. A protein of unknown function from *Bacteroides eggerthii* DSM 20697 with a lipocalin-like domain was revealed to be most phylogenetically similar to Scot1. One of the most common functions of the lipocalin-like domain is the transport of small molecules. An *E. coli* protein, YodA, with a lipocalin-like architecture is also known to bind metal ions to its barrel core. These results support the predicted structure-based annotation of Scot1 as a possible metal/small molecule carrier protein.

**Figure 6.**
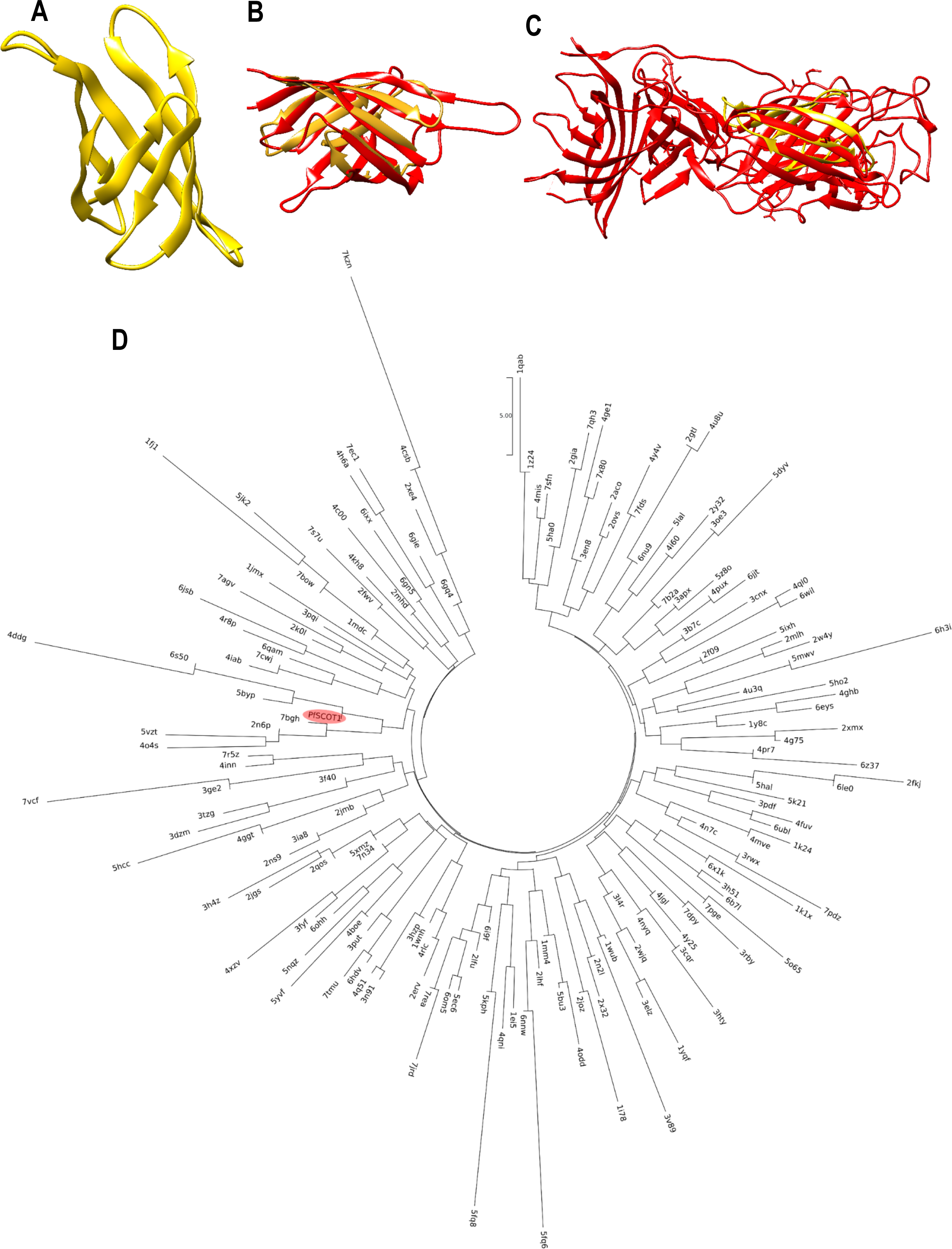
Bioinformatic analysis indicated that Scot1 is a metal/small molecule carrier protein. (**A**) The structure of PfScot1, as predicted using the Quark ab initio structure prediction web service, shows an open beta-barrel configuration. **(B)** Superposition of PfScot1 (gold) with chain M of PDB ID 2GTL (red) with a 0.9 Å RMSD value and **(C)** chain A of PDB ID 4O4X (red) with a 0.8 Å RMSD value. **(D)** Structure-based phylogenetic tree depicting the relative position of PfScot1 (highlighted) among its closest matching PDB counterparts listed in Table S1.

### The *Scot1* KO phenotype is not reversed by iron, zinc or cadmium

To determine whether Scot1 is a carrier of metal ions and whether supplementation with metals can rescue the KO phenotype. WT GFP and *Scot1* KO liver-stage cultures were grown in media supplemented with 50 µg /ml ferric ammonium citrate (FAC), 20 µM zinc chloride (ZnCl2) or 1µg/ml cadmium chloride (CdCl2). We observed the culture for the formation of detached cells at 65 hpi. We found detached cells in the WT GFP culture but not in the *Scot1* KO infected culture supplemented with metal ions (Figure 7A). Next, we checked the infectivity of detached cells from WT GFP and culture supernatant from *Scot1* KO parasites in Swiss mice. We found that culture treatment with metal ions did not affect detached cell formation or infection in mice (Table 3). This treatment also did not affect the maturation of EEFs or nuclear division (Figure 7B, and C). These data show that Scot1 is not a carrier of iron, zinc, or cadmium and can possibly act as a carrier for other small molecules. Another possibility is that Scot1 may detoxify metal/small molecules. However, this requires further investigation.

**Figure 7.**
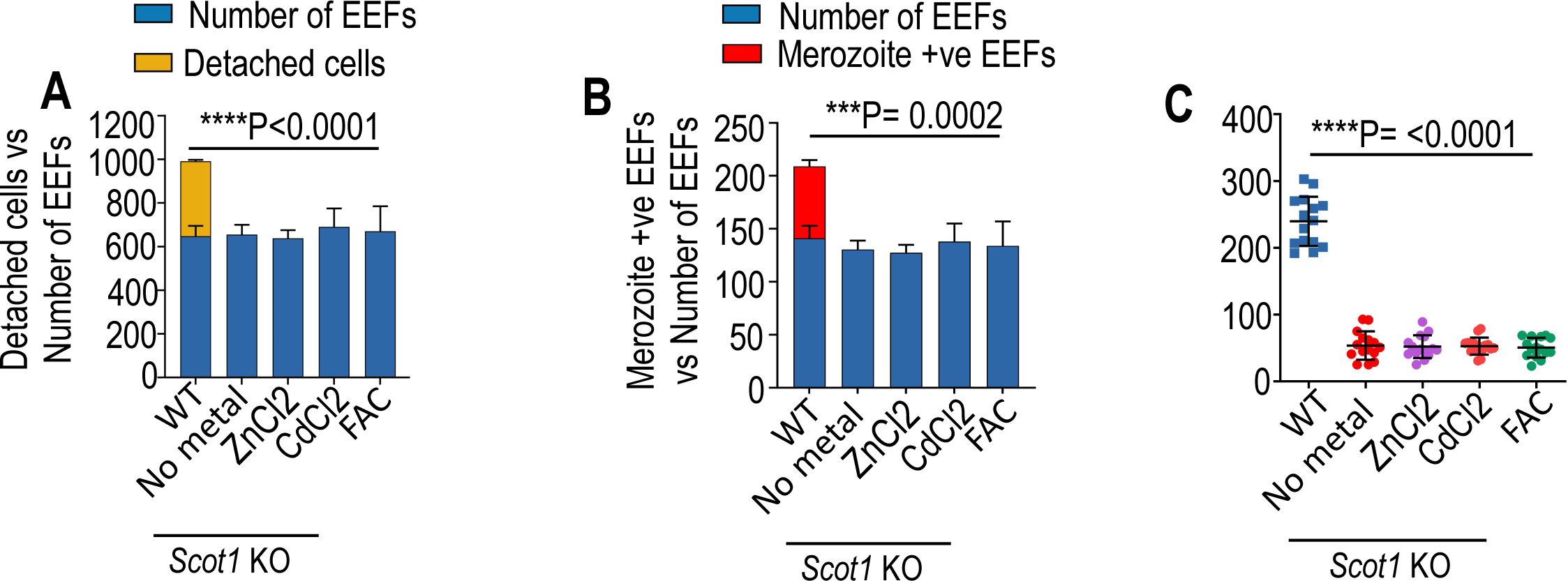
Metal ion supplementation does not rescue the *Scot1* KO phenotype. HepG2 cells were infected with either *Scot1* KO or WT GFP sporozoites and cultured in media supplemented with different concentrations of metal ions. The culture was observed at 65 hpi for the formation of detached cells and then fixed for IFA. (**A**) No detached cells were observed in the *Scot1* KO parasites supplemented with metal ions (****P<0.0001, one-way ANOVA). (**B**) Merozoites were not observed in the *Scot1* KO parasites supplemented with metal ions (***P= 0.0002, one-way ANOVA). (**C**) Impaired nuclear division in *Scot1* KO parasites supplemented with metal ions (****P<0.0001, one-way ANOVA). The data were obtained from two independent experiments performed in duplicate and are presented as the mean ± SEM.

**Table 3.** Infectivity of the merosomes in Swiss mice. Metal ion supplementation did not affect the *Scot1* KO phenotype. Blood smears were examined daily from day 1 onwards, and the mice were considered negative if parasites were not detected by day 20. NA, not applicable.

| Parasite | Number of detached cells injected | Mice positive/mice inoculated | Prepatent period (day) |
| --- | --- | --- | --- |
| WT | 10 | 10/10 | 2.5 |
| WT+ FAC | 10 | 10/10 | 2.6 |
| WT+ ZnCl<sub>2</sub> | 10 | 10/10 | 2.7 |
| WT+ CdCl<sub>2</sub> | 10 | 10/10 | 2.7 |
| *Scot1* KO | Supernatant | 0/10 | NA |
| *Scot1* KO + FAC | Supernatant | 0/10 | NA |
| *Scot1* KO + ZnCl<sub>2</sub> | Supernatant | 0/10 | NA |
| *Scot1* KO + CdCl<sub>2</sub> | Supernatant | 0/10 | NA |

### Immunization with *Scot1* KO sporozoites protects against malaria

Immunizations with sporozoites that arrest in the liver elicit a long-term protective host response and protect against infectious sporozoite challenge (27). We tested whether immunization with *Scot1* KO sporozoites confers protection from WT sporozoite infection. We observed that while all salivary gland debris-immunized mice developed blood-stage infection, the *Scot1* KO-immunized mice remained negative for the duration of the experiment (Table 4). These results are especially impressive since immunization of C57BL/6 mice with irradiated sporozoites rarely results in the immunity observed with *Scot1* KO sporozoites (28). Preerythrocytic protection after sporozoite immunization has revealed the induction of antibodies against CSP, suggesting the importance of humoral immunity (29, 30). We used IFA (Immunofluorescence assay), which recognizes both stages, to determine the serum reactivity against sporozoites and EEFs (Figure S7).

**Table 4.** Immunization of C57BL/6 mice with *Scot1*-KO sporozoites protects against infection. Mice were immunized with the indicated number of *Scot1* KO sporozoites and challenged on day 10 after the last immunization with *P. berghei* WT sporozoites. All immunized mice were protected from WT *P. berghei* sporozoite challenge. NA, not applicable.

| Experiment | Group | Immunization dose | Challenge day | Challenge dose | No. of patent/no. of challenged | Prepatent period (day) |
| --- | --- | --- | --- | --- | --- | --- |
| **1** | Control | (3x) SG debris | 10 | 5,000 | 5/5 | 3 |
| | *Scot1* KO | 1x) $5 \times 10^4$ and (2x) $2.5 \times 10^4$ | 10 | 5,000 | 0/5 | NA |
| **2** | Control | (3x) SG debris | 10 | 5,000 | 5/5 | 3 |
| | *Scot1* KO | 1x) $5 \times 10^4$ and (2x) $2.5 \times 10^4$ | 10 | 5,000 | 0/5 | NA |
| **3** | Control | (3x) SG debris | 10 | 5,000 | 5/5 | 3 |
| | *Scot1* KO | (3x) $2 \times 10^4$ | 10 | 5,000 | 0/5 | NA |

## Discussion

Sporozoites infect liver cells, where they mature into hepatic merozoites that invade red blood cells. Several sporozoite and liver stage-specific proteins have been implicated during liver stage development (5, 10, 31–35). Among them, P36 and P52 are microneme-localized proteins. Disruption of these genes led to growth arrest of the parasite in the liver and yielded GAP. Immunization with GAP parasites elicits immune responses that protect rodents and humans from an infectious sporozoite challenge (Dijk et al., 2005; Kublin et al., 2017; Labaied et al., 2007; A.-K. Mueller et al., 2005; Vaughan et al., 2009). In this study, we investigated the role of a sporozoite and liver stage-specific protein and showed that the expression of Scot1 is similar to the distribution pattern of micronemes. Our analysis of the subcellular localization of Scot1 and TRAP confirmed that Scot1 is a micronemal protein. After establishing the EEF, the micronemes are completely discarded into the PV (3). Parasites lacking the *Plasmodium* autophagy pathway protein Atg7 fail to expel the microneme into the PV (4). We found that Scot1 does not play a role in eliminating unneeded micronemes during liver stage development. This result indicated that the elimination of micronemes is regulated by the *Plasmodium* autophagy pathway (4) and that microneme-resident proteins are not involved in this process. Several micronemal proteins are involved in the hepatocyte invasion of sporozoites (5, 6, 8), whereas *Scot1* KO sporozoites invade hepatocytes normally, suggesting that Scot1 does not perform multiple essential functions and only plays a role in the development of the EEF. Like P52 and P36 mutants, *Scot1* KO parasites are arrested in the liver. However, compared with early-attenuating P52 and P36 mutants, *Scot1* KO parasites attenuate during late liver stage development (10).

A lack of Scot1 had no effect on blood or mosquito stage development. Scot1 was also found to be dispensable in a *P. falciparum* genetic screen (39). Micronemal proteins have been implicated in the gliding motility and invasion of sporozoites (5, 6, 8). Next, we analysed the invasion ability of *Scot1* KO sporozoites and found it to be normal. Furthermore, the early- to mid-liver-stage development of the *Scot1* KO parasites was normal, and the number and size of the EEFs were comparable to those of the WT GFP. In contrast, *Scot1* deletion affected late liver stage development. Earlier known GAPs, such as UIS3, UIS4 (upregulated in infectious sporozoites) and SPELD mutants, are arrested early before extensive DNA replication (31, 32, 34), while the type II fatty acid synthesis (FASII) pathway and RNA-binding protein *PlasMei2,* liver-stage antigen-1, and liver-specific protein 2 mutants are arrested late in liver stage development (36, 40–43).

The structural superposition with the SCOP and PDB databases using the FATCAT and DALI web services and subsequent superposition of the results in UCSF Chimera show that the model aligns with the haemoglobin linker chain L1 (chain M of PDB ID 2GTL (red)) with a 0.9 Å RMSD value and chain A of the crystal structure of the vaccine antigen Transferrin Binding Protein B (TbpB) (PDB ID: 4O4X (red)), a metal transport protein with a 0.8 Å RMSD value. The metal transport function prediction of Scot1 suggested that this could be another factor for its phenotype in the liver. The role of metals during liver stage development has been reported previously, and the role of a metal ion transporter, ZIPCO, was found to be critical for EEF development (44). Metals are associated with several processes in mammals and are implicated in oxygen transport in hemoglobin. Iron has been reported to play an essential role in DNA replication as a cofactor for ribonucleotide reductases (45) and metalloproteins. Iron is also essential for DNA replication as a cofactor of ribonucleotide reductases (45) and metalloproteins that contain an iron-sulfur (Fe-S) cluster (46). A study demonstrated that the ZIP protein Leishmania iron transporter 1 is essential for parasite replication in macrophage phagolysosomes (47, 48). The *Scot1* KO phenotype could not be reversed by supplementation with exogenously provided metal ions. A likely explanation for the developmental arrest of Scot1 in the liver stages may be that Scot1 is not a carrier of iron, zinc, or cadmium and can possibly act as a carrier for other small molecules. Another possibility is that Scot1 may detoxify metal/small molecules. In fact, a vacuolar iron transporter (VIT) has been described in *P. berghei* and is involved in detoxifying excess iron (49). Whether Scot1 plays a role in detoxifying excess metal ions requires further investigation.

Immunizations with genetically attenuated sporozoites that arrest in the liver elicit a long-term protective host response (50). We found that vaccination of C57BL/6 mice with *Scot1* KO sporozoites conferred complete protection against WT sporozoite challenge. Interestingly, late liver-arresting parasites provide superior protection compared to early-arresting parasites (51). Late liver-arresting parasites express late-stage antigens and a subset of antigens that are common to blood stages; however, whether Scot1 GAP exhibits superior protection needs further investigation. Despite the discovery of a growing list of GAPs, there have been some occasional breakthrough infections in a few GAPs. To overcome these limitations, double- or triple-attenuated GAPs can be generated by combination with known late-arresting GAPs (36, 40, 41), which increase the degree of attenuation of these parasites to prevent any possible breakthrough infection. Our investigations demonstrated that Scot1 is essential for late liver-stage development and could be used as a GAP vaccine.

## Materials and methods

### Parasites and mice

*P. berghei* ANKA (MRA 311) and *P. berghei* ANKA GFP (MRA 867 507 m6cl1) were obtained from BEI Resources, USA. Swiss albino and C57BL/6 mice were used for parasite infections. All animal procedures were approved by the Institutional Animal Ethics Committee at CSIR-Central Drug Research Institute, India (approval no: IAEC/2018/49).

### Amino acid sequence analysis

Interestingly, a study investigating the transcriptional landscape of *Plasmodium vivax* sporozoites revealed several upregulated transcripts with strong orthologues in the sporozoite stages of other species (52). We selected Scot1, which was the top-ranked gene in the transcriptome analysis. The *P. berghei* Scot1 (PBANKA_1411000) sequence was obtained from PlasmoDB (https://plasmodb.org/plasmo/app), and NCBI BLAST (https://blast.ncbi.nlm.nih.gov/Blast.cgi) was used to search for similar sequences in different organisms. Multiple sequence alignment (MSA) was performed using ClustalW. A sequence similarity matrix was prepared using the smith-waterman algorithm implemented in EMBOSS Water (https://www.ebi.ac.uk/jdispatcher/psa/emboss_water). The presence of the signal peptide and transmembrane domain was predicted using SignalP (https://services.healthtech.dtu.dk/services/SignalP-6.0) and TMHMM (https://services.healthtech.dtu.dk/services/TMHMM-2.0/) (53). A phylogenetic tree was constructed using MEGA11 software and the EMBL Interactive Tree Of Life (iTOL) service (https://itol.embl.de) (54).

### Generation of transgenic Scot1-3XHA-mCherry parasites to study their expression and localization

For the endogenous tagging of *Scot1* (PBANKA_1411000) with 3XHA-mCherry, two fragments, F1 (0.84 kb) and F2 (0.52 kb), were amplified using primers 1165/1171 and 1167/1168 and cloned into the pBC-3XHA-mCherry-hDHFR vector at *Xho*I/*Bgl*II and *Not*I/*Asc*I, respectively, as previously described (55). The targeting cassette was transfected into *P. berghei* ANKA schizonts (56), clonal lines were obtained by limiting dilution of the parasites, and correct integration was confirmed by diagnostic PCR using primers 1169/1218 and 1215/1170 (Table S2). Next, the mosquito cycle was initiated to observe the expression of mCherry in sporozoites and liver stages as previously described (55).

### Generation of *Scot1* knockout and complemented *P. berghei* lines

To delete *Scot*1, two fragments, F3 (0.63 kb) and F4 (0.52 bp), were amplified using the primer sets 1172/1173 and 1167/1168 and cloned into *Sal*I and *Not*I/*Asc*I in the pBC-GFP-hDHFR:yFCU vector. The targeting cassette was transfected into *P. berghei* schizonts, and clonal lines were obtained as described above. Site-specific 5’ and 3’ integrations were confirmed by diagnostic PCR using primers 2222/1225 and 1215/1170, respectively (Table S2). For complementation, fragment F5 was amplified from *P. berghei* genomic DNA using primers 1172/1168, transfected into *Scot1* KO schizonts, and selected using 5-fluorocytosine (MP Biomedicals, 219979201) as previously described (55). Restoration of the *Scot1* gene was confirmed using primers 1593/1594.

### Western blot analysis

Purified sporozoites were resuspended in Laemmli buffer (Bio-Rad, 1610747), analysed by SDS‒PAGE and transferred onto nitrocellulose membranes (Bio-Rad, 1620112), and the remaining procedures were performed as previously described (57). Briefly, the membrane was blocked with 1% BSA/PBS, followed by incubation with an anti-mCherry antibody developed in rabbits (diluted 1:500, Novus, NBP 2-25157) and an HRP-conjugated anti-rabbit secondary antibody (Amersham, NA934). The blot was developed using ECL Chemiluminescent Substrate (Bio-Rad, 1705060), and signals were detected using a ChemiDoc XRS+ System (Bio-Rad, USA). As a loading control, the blot was stripped and reprobed with an anti-TRAP antibody (diluted 1:200) (26).

### Sporozoite infectivity

To determine the in vivo infectivity of the *Scot1* KO sporozoites, C57BL/6 mice were either intravenously inoculated with salivary gland sporozoites or infected by mosquito bite, and the appearance of the parasite in the blood was observed by performing a Giemsa-stained blood smear. Another group of mice injected with 5,000 sporozoites was sacrificed at 40 and 55 hpi (hours post-infection), and the livers were harvested and homogenized in TRIzol reagent (Invitrogen, USA). To assess the invasion of the *Scot1* KO parasites, HepG2 cells were infected with sporozoites, and the culture was fixed at 1 hpi and immunostained before and after permeabilization with an anti-CSP monoclonal antibody (3D11) (30) as previously described (58). To observe EEF development, HepG2 cells were infected with salivary gland sporozoites and fixed at different time points using 4% paraformaldehyde as previously described (55). For the EEF and detached cell development assays, 5,000 and 40,000 sporozoites/well were added to 48-well and 24-well plates, respectively.

### Real-time PCR

Total RNA was isolated using TRIzol reagent, and cDNA was synthesized as previously described (59). The 18S rRNA copy number was determined by absolute quantification of gene-specific standards (60) using primers 1195/1196. The 18S rRNA copy number was normalized to that of mouse glyceraldehyde-3-phosphate dehydrogenase (GAPDH) using primers 1193/1194.

### Immunofluorescence assay

For the localization of Scot1 in sporozoites, purified salivary gland sporozoites were allowed to settle and dry on 12-well slides (Thermo Fisher Scientific, USA). Sporozoites were fixed using 4% paraformaldehyde (Sigma‒Aldrich, HT5012), permeabilized with 0.1% Triton-X-100 (Sigma‒Aldrich, T8787) for 10 min at RT and incubated with anti-mCherry developed in mouse (diluted 1:1000, Biolegend, 677702) and anti-TRAP (diluted 1:200, rabbit polyclonal) (26) antibodies for 2 h at RT. The mCherry and TRAP signals were revealed using Alexa Fluor 594-conjugated anti-mouse IgG and Alexa Fluor 488-conjugated anti-rabbit IgG, respectively (diluted 1:500; Invitrogen). The EEFs fixed at different time points were permeabilized using methanol, blocked using 1% BSA/PBS and incubated with primary antibodies as previously described (57). To visualize the microneme elimination pattern, infected cultures fixed at different time points were immunostained with an anti-TRAP antibody (diluted 1:200) (26). The primary antibodies used were anti-mCherry (diluted 1:500), anti-UIS4 (diluted 1:1,000, rabbit polyclonal) (31), anti-TRAP, anti-MSP1 (diluted 1:5000, mouse monoclonal) (61), and anti-ACP (diluted 1:1,000, rabbit polyclonal) (62). The signals were revealed using Alexa Fluor 488-conjugated or Alexa Fluor 594-conjugated secondary antibodies (diluted 1:500; Invitrogen). Nuclei were stained with Hoechst 33342 (Sigma‒Aldrich, 41399), and the coverslips were mounted using Prolong Diamond antifade reagent (Invitrogen, P36970). Representative images were acquired using FV1000 software on a confocal laser scanning microscope (Olympus BX61WI) using UPlanSAPO 100x (NA 1.4, oil) or 63x (NA 0.25, oil).

### Bioinformatic analysis of Scot1 proteins

The 3D structure of PfScot1 was predicted using the Quark (https://zhanggroup.org/QUARK) ab initio structure prediction web service. Structure-based phylogenetic analysis was performed using the predicted structure of PfScot1. 3D Phylofold is a recently developed method used to repurpose existing antivirals for COVID-19 treatment (63). The DALI webserver was used to identify structural matches of PfScot1 in the PDB database (64). The top hits from the results were analysed using 3Dphylofold to generate a structure-based similarity matrix. The matrix was further analysed using Mega to generate a nearest-neighbor dendrogram (https://www.megasoftware.net).

### Effects of metal supplementation on EEF maturation

HepG2 cells were infected with WT GFP or *Scot1* KO sporozoites as described above. Two hpi, fresh media containing 50 µg /ml ferric ammonium citrate (FAC), 20 µM zinc chloride (ZnCl2) or 1µg/ml cadmium chloride (CdCl2) were added. The media was changed every 12 hours, supplemented with metals. At 65 hpi, culture was observed for the formation of detached cells and then fixed with 4% paraformaldehyde (Sigma‒Aldrich, HT5012). The EEFs were stained with an anti-MSP1 antibody, and the nuclei were stained with Hoechst 33342 (Sigma‒Aldrich, 41399) as described above. Coverslips were mounted using Prolong Diamond antifade reagent (Invitrogen, P36970). Images were acquired using FV1000 software on a confocal laser scanning microscope (Olympus BX61WI) using UPlanSAPO 100x (NA 1.4, oil) or 63x (NA 0.25, oil).

### Immunization and challenge experiments

Six- to eight-week-old female C57BL/6 mice were primed intravenously with 50,000 salivary gland sporozoites and boosted twice with 25,000 at an interval of two weeks. The salivary gland debris of uninfected mosquitoes was injected into the control group. Another group was immunized thrice with 20,000 sporozoites. The control and immunized groups were challenged with 5,000 infectious WT sporozoites 10 days after the last immunization. Parasitemia was monitored daily by making Giemsa-stained blood smears.

### Immunized mouse serum reactivity

To visualize the reactivity of sera obtained from the immunized mice, sporozoites and EEFs were stained with pooled serum (diluted 1:50), and signals were revealed using Alexa Fluor 594-conjugated anti-mouse IgG (diluted 1:500; Invitrogen). To identify the sporozoites and EEFs, anti-TRAP (26) and anti-UIS4 antibodies (31), respectively, were used. Nuclei were stained with Hoechst 33342.

### Statistical analysis

The data are presented as the mean ± SEM or mean ± SD. Statistical analysis was performed using GraphPad Prism 9 software. As indicated, two-tailed, unpaired Student’s t test and one-way ANOVA were used to determine the statistical significance.

### Availability of data and material

All the data are available within this manuscript, and the raw data are available from the corresponding author upon reasonable request. Materials generated in this study are available from the corresponding author upon request.

## Acknowledgments

We thank Dr. Robert Menard (Institute Pasteur, France) for the pBC-GFP-hDHFR vector, which was modified by Dr. P.N. Srivastava. We thank Dr. Kota Arun Kumar (University of Hyderabad, India) for the pBC-3XHA-mCherry-hDHFR vector. We thank Dr. Anthony A. Holder (The Francis Crick Institute, UK), Drs. Photini Sinnis and Sean Prigge (Johns Hopkins University, USA) for anti-MSP1, anti-UIS4, and anti-ACP antibodies, respectively. AG, Nirdosh, RD, AM, SKN and PNS were supported by CSIR, UGC ICMR and DBT Research Fellowships. We acknowledge the THUNDER (BSC0102) and MOES (GAP0118) Intravital and Confocal Microscopy Facilities of CSIR-CDRI. The graphics were created with Biorender.com. This work was supported by a Science and Engineering Research Board grant (EMR/2016/006487).

## Author contributions

SM conceived the idea, designed the experiments, analysed the data and wrote the manuscript. AG, RD, AM, SKN, N and PNS performed the experiments. All the authors have read and approved the manuscript.

## Supplemental Material

**Figure S1.** Phylogenetic tree.

**Figure S2.** Scot1 amino acid sequence analysis.

**Figure S3.** Generation of Scot1-3XHA-mCherry parasites and their development in the mosquito and liver stages.

**Figure S4.** Generation of *Scot1* KO and complemented parasites.

**Figure S5.** The development of *Scot1* KO parasites in mosquitoes.

**Figure S6.** *Scot1* KO sporozoites invade hepatocytes normally.

**Figure S7.** Immune sera recognize sporozoites and the EEF.

**Table S1.** Closest structure-based phylogenetic matches of PfSCOT1 from the PDB database.

**Table S2.** List of primers used in this study.

